# Kif17 phosphorylation regulates its ciliary localization and photoreceptor outer segment turnover

**DOI:** 10.1101/351122

**Authors:** Tylor R. Lewis, Sean R. Kundinger, Brian A. Link, Christine Insinna, Joseph C. Besharse

**Author notes:** Corresponding author (JCB). **Current addresses of TRL:** Department of Ophthalmology, Albert Eye Research Institute, Durham, NC 27710, USA.

## Abstract

**Background:** KIF17, a kinesin-2 motor that functions in intraflagellar transport, can regulate the onset of photoreceptor outer segment development. However, the function of KIF17 in a mature photoreceptor remains unclear. Additionally, the ciliary localization of KIF17 is regulated by a C-terminal consensus sequence (KRKK) that is immediately adjacent to a conserved residue (mouse S1029/zebrafish S815) previously shown to be phosphorylated by CaMKII. Yet, whether this phosphorylation can regulate the localization, and thus function, of KIF17 in ciliary photoreceptors remains unknown.

**Results:** Using transgenic expression in both mammalian cells and zebrafish photoreceptors, we show that phospho-mimetic KIF17 has enhanced localization to cilia. Importantly, expression of phospho-mimetic KIF17 is associated with greatly enhanced turnover of the photoreceptor outer segment through disc shedding in a cell-autonomous manner, while genetic mutants of *kif17* in zebrafish and mice have diminished disc shedding. Lastly, cone expression of constitutively active tCaMKII leads to a *kif17*-dependent increase in disc shedding.

**Conclusions:** Taken together, our data support a model in which phosphorylation of KIF17 promotes its ciliary localization. In cone photoreceptor outer segments, this promotes disc shedding, a process essential for photoreceptor maintenance and homeostasis. While disc shedding has been predominantly studied in the context of the mechanisms underlying phagocytosis of outer segments by the retinal pigment epithelium, this work implicates photoreceptor-derived signaling in the underlying mechanisms of disc shedding.

## Background

Cilia, organelles that project from the cell body and contain a microtubule-based axoneme, have numerous roles in development, signal transduction, and cell motility [1] and have garnered widespread interest due to associations with a widespread set of disorders termed ciliopathies [2]. Although the ciliary membrane is continuous with the plasma membrane, the transition zone at the base of the cilium is believed to function as a ciliary gate involved in regulating protein entry and exit [3]. While there has been conflicting evidence regarding the permeability barrier at the base of cilia [4, 5], it has been shown that ciliary entry of the kinesin-2 family member, KIF17, is dependent on a putative nuclear localization signal (NLS) **(Figure 1A)** that can additionally function as a ciliary localization signal (CLS) [6, 7]. Although KIF17 has been predominantly studied as a ciliary or dendritic motor [8], it has been reported to localize to the nucleus and function in transcriptional regulation [9, 10]. Yet the context under which a single conserved sequence can function as either a NLS or CLS remains unclear. Calcium/calmodulin-dependent protein kinase II, or CaMKII, has been shown to phosphorylate the C-terminus of KIF17 to regulate binding and release of Mint1, a scaffolding protein for motor cargo, in neurons [11]. Interestingly, this conserved phosphorylation site lies immediately adjacent to the NLS/CLS in KIF17 **(Figure 1A)**.

**Figure 1.**
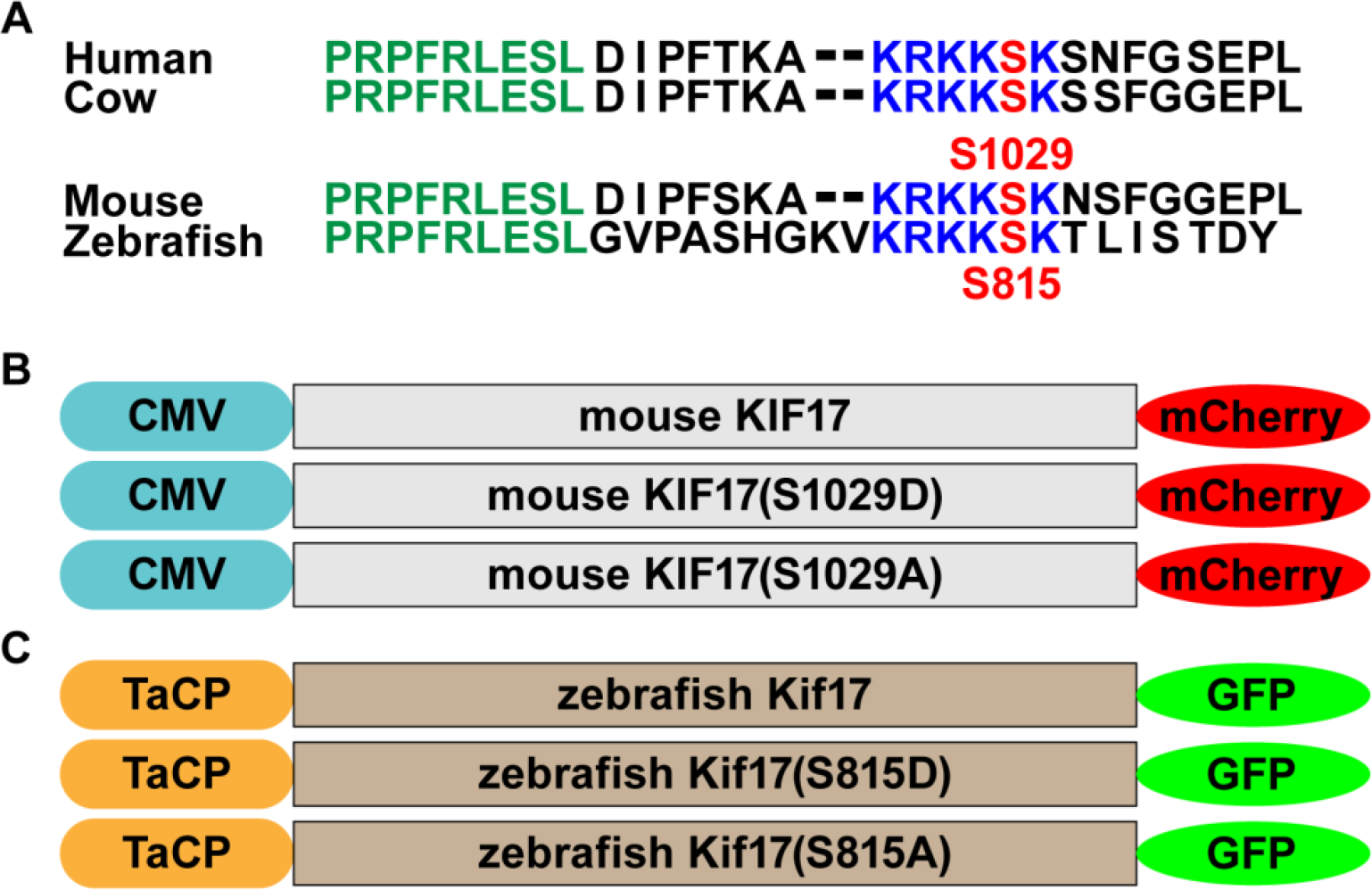
Schematic of transgenic phospho-mutant KIF17 constructs. **(A)** The amino acid sequence of the C-terminus of KIF17 in various species is depicted. Homologous sequences are shown in color (green, blue, and red). The green conserved sequence has been shown to interact with the IFT-B complex [7], while the blue conserved sequence functions as a NLS or CLS [6]. The conserved serine (red), mouse S1029/zebrafish S815, has been shown to be phosphorylated by CaMKII [11]. **(B)** For mammalian cell culture experiments, three different mCherry-tagged constructs of mouse KIF17 under control of the ubiquitous CMV promoter were generated: wild-type KIF17, phospho-mimetic KIF17(S1029D), and phospho-deficient KIF17(S1029A). **(C)** For zebrafish cone photoreceptor experiments, three different GFP-tagged constructs of zebrafish under control of the cone-specific transducin-α promoter (TaCP) were generated: wild-type Kif17, phospho-mimetic Kif17(S815D), and phospho-deficient Kif17(S815A).

KIF17 is expressed in photoreceptors [12], which have a modified primary cilium called the outer segment (OS) that contains the components required for phototransduction [13]. Although KIF17 may regulate the onset of OS morphogenesis [14], there has been no evidence of a role for KIF17 in a mature photoreceptor [15]. As KIF17 can accumulate at the ciliary tip of photoreceptors [16], like mammalian cell lines [6, 7, 15, 17], it is possible that KIF17 may have some role in distal tip maintenance. Interestingly, the distal tip of the OS is phagocytized by the opposing retinal pigment epithelium (RPE) daily in a process called disc shedding [18]. The molecular mechanisms underlying disc shedding have been predominantly studied at the level of the RPE, but exposed phosphatidylserine (PS) on the tips of OS precedes disc shedding [19], suggesting photoreceptor signaling contributes to the underlying mechanism of disc shedding.

In this work, we express three different forms of mCherry-tagged mouse KIF17 (wild-type, phospho-mimetic S1029D, and phospho-deficient S1029A) **(Figure 1B)** in four separate mammalian cell lines and show that phospho-mimetic KIF17(S1029D) accumulates extensively in cilia compared to either wild-type KIF17 or phospho-deficient KIF17(S1029A). Significant differences in nuclear localization of the transgenes were only seen in two of four cell lines, where KIF17(S1029D) had restricted nuclear localization. Additionally, with cone expression of three different forms of GFP-tagged zebrafish Kif17 (wild-type, phospho-mimetic S815D, and phospho-deficient S815A) **(Figure 1C)**, we show that the accumulation of Kif17 in the zebrafish OS is controlled through this phosphorylation site. Cone expression of phospho-mimetic Kif17 leads to a three-fold increase in disc shedding, while the genetic mutant *kif17*^*mw405*^ has diminished disc shedding. Additionally, cone expression of a constitutively active CaMKII leads to a two-fold increase in disc shedding in wild-type, but not *kif17*^*mw405*^ mutant fish. Taken together, this work supports a model in which phosphorylation of Kif17 regulates its accumulation in cilia. In cone photoreceptors, phosphorylated Kif17 in the OS participates in a cell-autonomous process to promote disc shedding.

## Results

### Phospho-mimetic mouse KIF17 has enhanced ciliary localization

Published data suggest that the conserved KRKK sequence in the C-terminus of KIF17 can function as both a NLS and CLS [6]. Interestingly, a conserved phosphorylation site (S1029 in mice and S815 in zebrafish) lies immediately adjacent to this NLS/CLS **(Figure 1A)** and has been shown to be phosphorylated by CaMKII in neurons [11]. To further investigate this phosphorylation site with regards to ciliary and nuclear localization, we generated three different constructs of mouse KIF17 for mammalian cell expression with the CMV promoter: KIF17-mCherry, phospho-mimetic KIF17(S1029D)-mCherry, and phospho-deficient KIF17(S1029A)-mCherry **(Figure 1B)**. To investigate the extent to which this phosphorylation site controls ciliary and nuclear localization, we used four different ciliated mammalian cell lines: LLC-PK1, a pig kidney epithelial line; HEK-293, a human kidney line; hTERT-RPE1, a human retinal pigment epithelial line; and IMCD3, a mouse kidney line. With transient transfection of serum-starved cells, we can determine the extent of ciliary localization of each transgene with co-labeling of acetylated α-tubulin, which labels the axoneme, or nuclear localization with co-labeling of Hoechst **(Figure 2A)**. The frequency of ciliary localization of the phospho-mimetic KIF17(S1029D)-mCherry was significantly increased between two- to four-fold over the phospho-deficient KIF17(S1029A)-mCherry in all cell lines studied **(Figure 2B)**. Interestingly, the ciliary localization of wild-type KIF17-mCherry was never significantly different from the phospho-deficient KIF17(S1029A)-mCherry with both having a baseline level of ciliary localization, suggesting that phosphorylation enhances ciliary localization but does not absolutely control ciliary entry.

**Figure 2.**
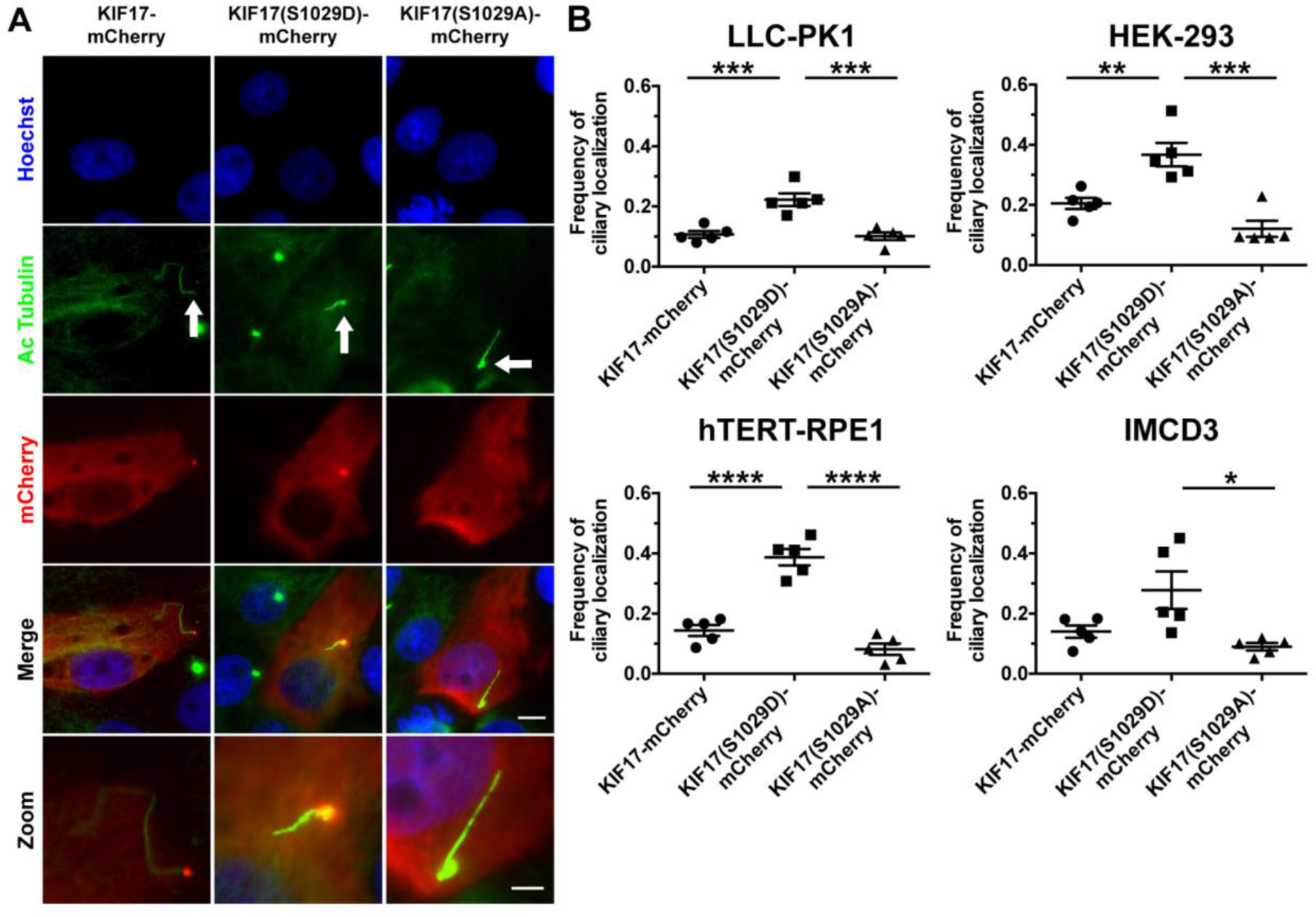
Phospho-mutations of S1029 regulate ciliary localization of KIF17. **(A)** LLC-PK1 cells were serum-starved to induce ciliogenesis and transfected with either KIF17-mCherry (top), KIF17(S1029D)-mCherry (middle), or KIF17(S1029A)-mCherry (bottom). 24 hours following transfection, cells were stained with Hoechst (blue) to label nuclei and acetylated a - tubulin (green) to label axonemes. The arrows mark the cilia. The zoom panel depicts a high-magnification of the cilium. Of note, transgenic KIF17 accumulates within the cilia (arrows), presumably at the ciliary tip as observed previously [6] [7] [15]. Cilia are often twisted and nonlinear. Additionally, there is generally significantly less nuclear than cytoplasmic localization of each KIF17 transgene. Scale bar is 10 μm for normal magnification images. Scale bar is 4 μm for zoom panel. **(B)** Quantification of frequency of ciliary localization of each transgene 24 hours posttransfection in various mammalian ciliated cell lines. In LLC-PK1 cells, ciliary localization of KIF17-mCherry (n=5 transfections, 710 cells), KIF17(S1029D)-mCherry (n=5, 859 cells), and KIF17(S1029A)-mCherry (n=5, 664 cells) is shown. In HEK-293 cells, ciliary localization of KIF17-mCherry (n=5 transfections, 690 cells), KIF17(S1029D)-mCherry (n=5, 1149 cells), and KIF17(S1029A)-mCherry (n=5, 886 cells) is shown. In hTERT-RPE1 cells, ciliary localization of KIF17-mCherry (n=5 transfections, 280 cells), KIF17(S1029D)-mCherry (n=5, 307 cells), and KIF17(S1029A)-mCherry (n=5, 279 cells) is shown. In IMCD3 cells, ciliary localization of KIF17-mCherry (n=5 transfections, 665 cells), KIF17(S1029D)-mCherry (n=5, 815 cells), and KIF17(S1029A)-mCherry (n=5, 845 cells) is shown.

To analyze nuclear localization, we calculated the ratio of average mCherry signal intensity between the nucleus and cytoplasm **(Figure S1)**. In two of the four cell lines (LLC-PK1 and HEK-293), there were no effects on transgene localization between the nucleus and cytoplasm despite there being a ciliary localization difference. However, in hTERT-RPE1 and IMCD3 cells, the phospho-mimetic KIF17(S1029D)-mCherry had a decreased nuclear to cytoplasmic ratio, suggesting inhibition of nuclear localization as compared to either wild-type KIF17-mCherry or phospho-deficient KIF17(S1029A)-mCherry. Yet, only in hTERT-RPE1 cells did the nuclear to cytoplasmic ratio exceed one. Taken together, these data support a model in which phosphorylation of the conserved S1029 residue of mouse KIF17 strongly promotes ciliary localization in four different cell lines. In contrast, nuclear localization of KIF17 may be negatively regulated by phosphorylation depending on cell type.

### Phospho-mimetic zebrafish Kif17 has enhanced OS localization

While phosphorylation of KIF17 may regulate ciliary localization in mammalian cells, our main interest was in investigating the effect of KIF17 phosphorylation in ciliary photoreceptor OS *in vivo*. We generated comparable zebrafish Kif17 constructs specifically for cone photoreceptor expression with the cone-specific transducin-α (*gnat2*) promoter, also known as TaCP: Kif17-GFP, phospho-mimetic Kif17(S815D)-GFP, and phospho-deficient Kif17(S815A)-GFP **(Figure 1C)**. While previous work shows that Kif17-GFP can accumulate at the distal tip of the OS [16], we used transient injections of each of these constructs into wild-type zebrafish embryos to quantify localization within the OS of two sub-types of zebrafish cones: blue and green absorbing cones. Supporting the mammalian cell work, phospho-mimetic Kif17(S815D)-GFP accumulated in the OS, while phospho-deficient Kif17(S815A)-GFP appeared to accumulate at the base of the OS **(Figures 3A, S2)**. The accumulation of phospho-mimetic Kif17(S815D)-GFP in the OS was sevenfold greater than that of phospho-deficient Kif17(S815A)-GFP **(Figure 3B)**. Additionally, line intensity analysis reveals that phopsho-mimetic Kif17(S815D)-GFP localizes evenly along the entire length of the OS, while phospho-deficient Kif17(S815A)-GFP exhibited a specific accumulation at the base of the OS **(Figure 3C)**. We did not observe any evidence of nuclear localization of the zebrafish Kif17-GFP constructs **(Figure S2)**. These data further support a model in which phosphorylation of Kif17 leads to an enhanced localization in the ciliary photoreceptor OS.

**Figure 3.**
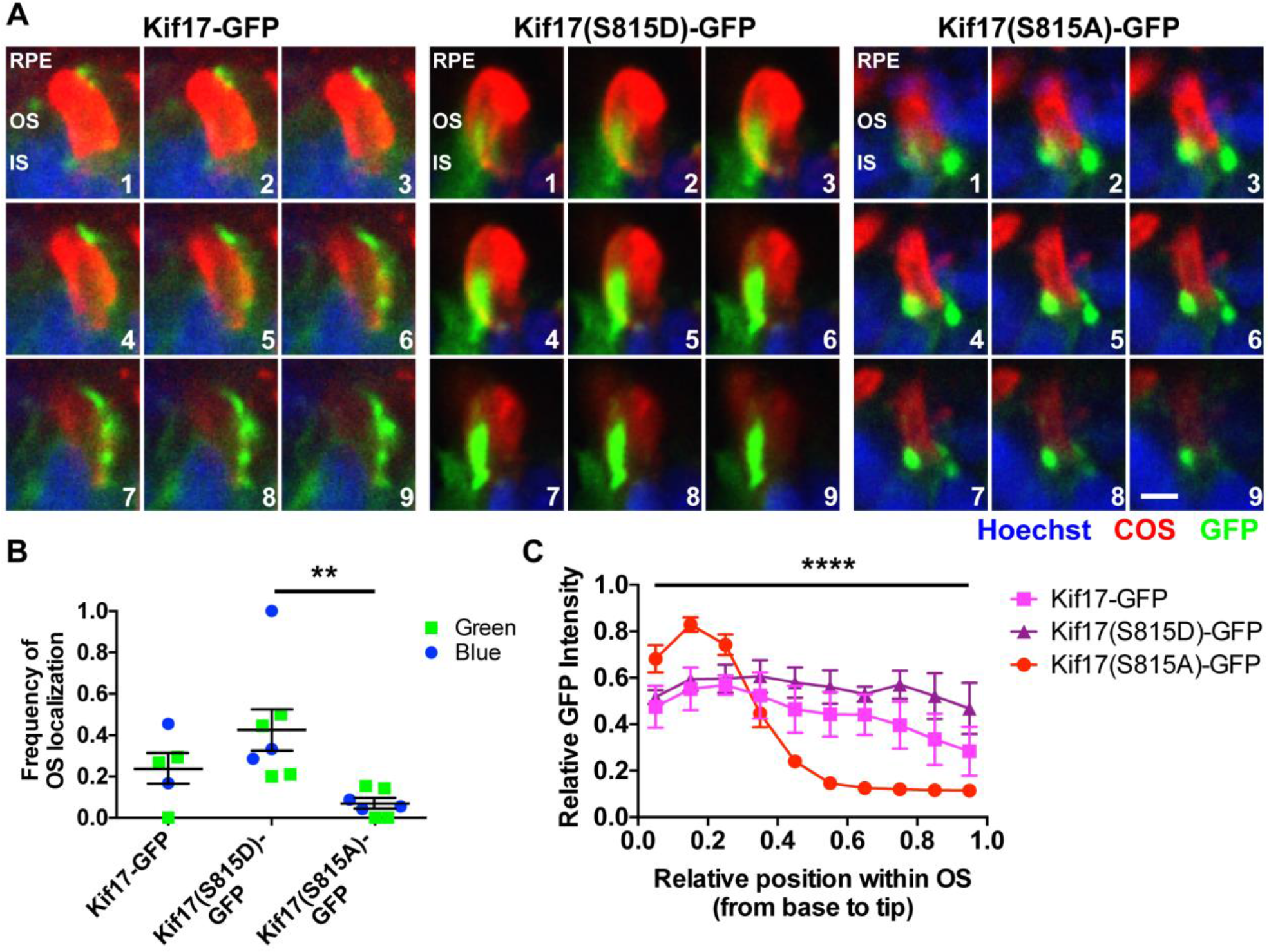
Phospho-mutations of S815 regulate photoreceptor OS localization of Kif17. **(A)** 5 dpf larvae previously injected at the one cell stage with one of three different transgenic constructs under control of the TaCP promoter for expression in cone photoreceptors: Kif17-GFP (left), phospho-mimetic Kif17(S815D)-GFP (middle), and phospho-deficient Kif17(S815A)-GFP (right) were stained with Hoechst (blue) to label nuclei and blue cone opsin (red) to label cone OS. A z-series of confocal images was taken with 0.2 μm steps through the depth of the photoreceptor OS. Shown are montages of sequential images in the z-series revealing that Kif17(S815D)-GFP exhibits significant accumulation in the OS while Kif17(S815A)-GFP accumulates at the base of the OS. Note that in cone photoreceptors, the ciliary axoneme extends along the side of the opsin-containing discs. Scale bar is 1 μm. **(B)** Quantification of the frequency of OS accumulation of Kif17-GFP (n=5 total injected larvae: 3 stained for green opsin, 2 stained for blue opsin; 60 cells),Kif17(S815D)-GFP (n=7: 4 green, 3 blue; 62 cells), and Kif17(S815A)-GFP (n=7: 4 green, 3 blue; 126 cells) when co-labeled with either green opsin (square) or blue opsin (circle). **(C)** Line intensity analysis of the GFP signal for Kif17-GFP (pink line, n=5 total injected larvae: 3 stained for green opsin, 2 stained for blue opsin; 17 cells), Kif17(S815D)-GFP (purple line, n=5: 3 green, 2 blue; 17 cells), and Kif17(S815A)-GFP (red line, n=5: 3 green, 2 blue; 25 cells). Two-way ANOVA was performed to determine significance in the different interaction between GFP signal intensity pattern and transgene (p<0.0001, ****).

### KIF17 promotes cell-autonomous disc shedding

While we have previously shown a developmental role of KIF17 in regulating the onset of OS morphogenesis [14], there is no evidence of a role of KIF17 in mature photoreceptors [15]. We hypothesized that investigating the temporal expression patterns of *Kif17* in mature photoreceptors throughout the diurnal light:dark cycle could shed light on a potential function of KIF17. We performed qPCR on mouse retinas that were collected every four hours during a 24 hour period beginning at light onset, indicated as Zeitgeber Time (ZT) 0, through dark onset (ZT 12), and stopping at ZT 20, or 20 hours following light onset. Interestingly, there is a single peak in *Kif17* expression in the mouse retina at ZT 4, just following light onset, which subsequently returns to a basal level **(Figure S3A)**. We also performed qPCR on 14 dpf zebrafish eyes that were collected at nine discrete timepoints over a 24 hour period beginning at light onset. In contrast to the mice, which are kept on a 12 hour:12 hour light:dark cycle, the zebrafish are maintained on a 14 hour:10 hour light:dark cycle, so that ZT 14 indicates dark onset. While there is an increase in zebrafish *kif17* expression immediately after light onset (ZT 1.5), there is a second, more intense peak just following dark onset (ZT 16 and ZT 18) **(Figure S3B)**. While *Kif17* appears to be rhythmically expressed in both mouse and zebrafish, mice have a single peak of expression associated with light onset, whereas zebrafish have two peaks of expression associated with both light and dark onset.

Due to the association of *kif17* expression with both light and dark onset in zebrafish, we hypothesized that Kif17 could be involved in disc shedding, the daily process of OS turnover in which RPE cells adjacent to the OS phagocytize and degrade shed OS tips. To analyze disc shedding, we performed transmission electron microscopy (TEM) on 5dpf zebrafish larvae that had been injected with the cone-expressing plasmids encoding either one of the three Kif17 transgenes or a soluble GFP control. Lacking transposase, the plasmids do not incorporate into the genome but rather facilitate episomal expression of transgenes. Interestingly, transient expression of the phospho-mimetic Kif17(S815D)-GFP was sufficient to drive an increase in the number of phagosomes compared to wild-type Kif17, phospho-deficient Kif17, or the soluble GFP control **(Figure S4)**. Because of the potential for interactions of the Kif17 transgenes with endogenous wild-type Kif17 and the transient and mosaic nature of expression of injected plasmids, we generated stable lines of each of our cone-expressing transgenes (Kif17-GFP, phospho-mimetic Kif17(S815D)-GFP, and phospho-deficient Kif17(S815A)-GFP) on a mutant background called *kif17*^*mw405*^ lacking endogenous Kif17 [14]. Following several generations of outcrossing to ensure only a single copy was inserted for each transgene **(Figure S5A,B)**, we collected 7dpf larvae with comparable levels of expression **(Figure S5A,C)** at ZT 1.5 and performed TEM to analyze phagosomes **(Figure 4A)**. There was a dramatic three-fold increase in the number of phagosomes observed in the RPE in retinas with cone expression of Kif17(S815D)-GFP as compared to either Kif17-GFP or Kif17(S815A)-GFP **(Figure 4B)**, while the average size of phagosomes was unchanged **(Figure 4C)**. Surprisingly, while there was a significant increase in disc shedding associated with Kif17(S815D)-GFP expression in cones, there was no decrease in average OS area **(Figure 4D)**, suggesting that the excess in disc shedding may compensate for excess OS morphogenesis. However, an upregulation of opsin mRNA expression, which encodes the major protein of the OS, was not observed **(Figure S6)**.

**Figure 4.**
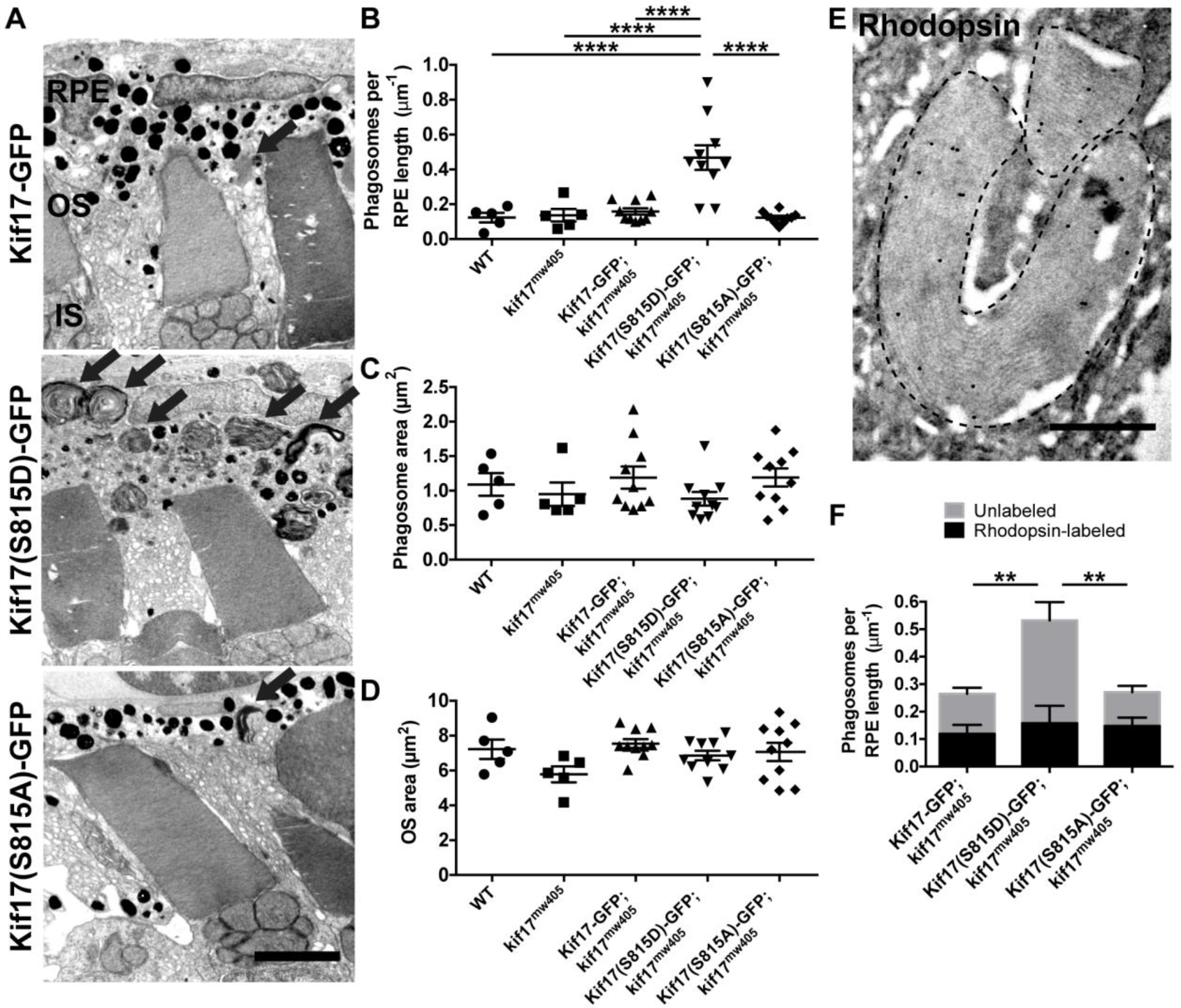
Phospho-mimetic Kif17(S815D) increases disc shedding. **(A)** TEM images of 7dpf *kif17*^mw405^ larvae that are stably expressing either Kif17-GFP (left), phospho-mimetic Kif17(S815D)-GFP (middle), or phospho-deficient Kif17(S815A)-GFP (right) in cones collected at ZT 1.5. Black arrows indicate phagosomes. Scale bar is 2 μm. IS is inner segment. **(B)** Quantification of the number of phagosomes for wild-type (n=5 larvae, 1273 μm of RPE); *kif17*^mw405^ (n=5, 1170 |jm of RPE); Kif17-GFP, *kif17*^*mw405*^ (n=10, 2677 μm of RPE); Kif17(S815D)-GFP, *kif17*^*mw405*^ (n=10, 2938 μm of RPE); and Kif17(S815A)-GFP, *kif17*^*mw405*^ (n=10, 2875 μm of RPE) larvae. **(C)** Quantification of the area of phagosomes for wild-type (n=5 larvae, 403 phagosomes); *kif17*^*mw405*^ (n=5, 392 phagosomes); Kif17-GFP, *kif17*^*mw405*^ (n=10, 530phagosomes); Kif17(S815D)-GFP, *kif17*^*mw405*^ (n=10, 1601 phagosomes); and Kif17(S815A)-GFP, *kif17*^*mw405*^ (n=10, 547 phagosomes) larvae. **(D)** Quantification of OS size for wild-type (n=5 larvae, 268 OS); *kif17*^*mw405*^ (n=5, 255 OS); Kif17-GFP, *kif17*^*mw405*^ (n=10, 615 OS); Kif17(S815D)-GFP, *kif17*^*mw405*^ (n=10, 566 OS); and Kif17(S815A)-GFP, *kif17*^*mw405*^ (n=10, 605 OS) larvae. **(E)** Immunogold labeling of a 7 dpf TaCP:Kif17-GFP zebrafish retina with K62-171c, an antibody against bovine rhodopsin showing a positive labeling of a phagosome, as also depicted in Figure S7. Scale bar is 0.5 μm. **(F)** Analysis of immunogold labeling of rhodopsin-containing or unlabeled phagosomes in 7 dpf Kif17-GFP, *kif17*^*mw405*^ (n=5 larvae, 1031 |jm of RPE); Kif17(S815D)-GFP, *kif17*^*mw405*^ (n=5, 1268 μm of RPE); and Kif17(S815A)-GFP, *kif17*^*mw405*^ (n=5, 889 μm of RPE) larvae. Two-way ANOVA was performed to determine significance in the interaction between the transgene expressed and the concentration of rod or cone phagosomes (p=0.0264, *) as well as significance in the total number of phagosomes between transgenes (p=0.0071, **). Bonferroni’s multiple comparisons post-hoc test was performed to determine significance in the differences of unlabeled phagosomes between Kif17-GFP and Kif17(S815D)-GFP (p=0.0033, **) and between Kif17(S815A)-GFP and Kif17(S815D)-GFP (p=0.0013, **). There were no differences in numbers of rhodopsin-labeled phagosomes among the transgenes.

Because our TEM analysis does not distinguish between phagosomes from either cone or rod photoreceptors, we performed immunogold labeling of these transgenic larvae. Using a monoclonal antibody against bovine rhodopsin, K62-171c [20], that also specifically labels zebrafish rhodopsin-containing OS and phagosomes **(Figures 4E, S7)**, we show that cone expression of Kif17(S815D)-GFP leads to a three-fold increase in unlabeled phagosomes, presumably from cones, compared to larvae with cone expression of Kif17(S815A)-GFP **(Figure 4F)**. However, there is no significant change in the number of rhodopsin-labeled phagosomes. Additionally, we specifically labeled cone-derived phagosomes using an antibody against gnat2, the cone-specific transducin-α whose promoter was used to drive expression of the transgenes in this study. Although the immunogold labeling of cone transducin-α labeled both cone OS and phagosomes **(Figure S8A)**, the labeling efficiency was significantly lower than that of the rhodopsin antibody, presumably due to a lower concentration of gnat2 in the OS. Nonetheless, we show that cone expression of Kif17(S815D)-GFP leads to a two-fold increase in cone transducin-α-labeled phagosomes with no significant change in unlabeled phagosomes, presumably from rods **(Figure S8B)**. Taken together, these data suggest phosphorylation of Kif17 promotes both OS localization and subsequent cone disc shedding in a cell-autonomous manner.

### Loss of KIF17 diminishes disc shedding in zebrafish and mice

While transgenic cone expression of phospho-mimetic Kif17(S815D)-GFP promotes disc shedding, we sought to further investigate how the loss of *kif17* affects disc shedding. While we did not observe defects in disc shedding in *kif17*^*mw405*^ mutants at the single timepoint (ZT 1.5) analyzed in our experiments with the transgenic fish **(Figure 4B)**, we proposed that deficiencies of disc shedding in the *kif17*^*mw405*^ zebrafish would be better observed when analyzed throughout the entire day. We collected larvae at 14 dpf, when OS have reached their mature size [14], at the nine timepoints over a 24 hour period and performed TEM to image phagosomes in both wild-type and *kif17*^*mw405*^ zebrafish. We find that zebrafish disc shedding is highly rhythmic, with two peaks associated with both light and dark onset. Additionally, there is a significant decrease in the number of phagosomes shed in zebrafish lacking Kif17 (*kif17*^*mw405*^) as compared to controls **(Figure 5A)**. Although there were changes in the disc shedding rates between genotypes and throughout the day, there was no change in phagosome size **(Figure 5B)**.

**Figure 5.**
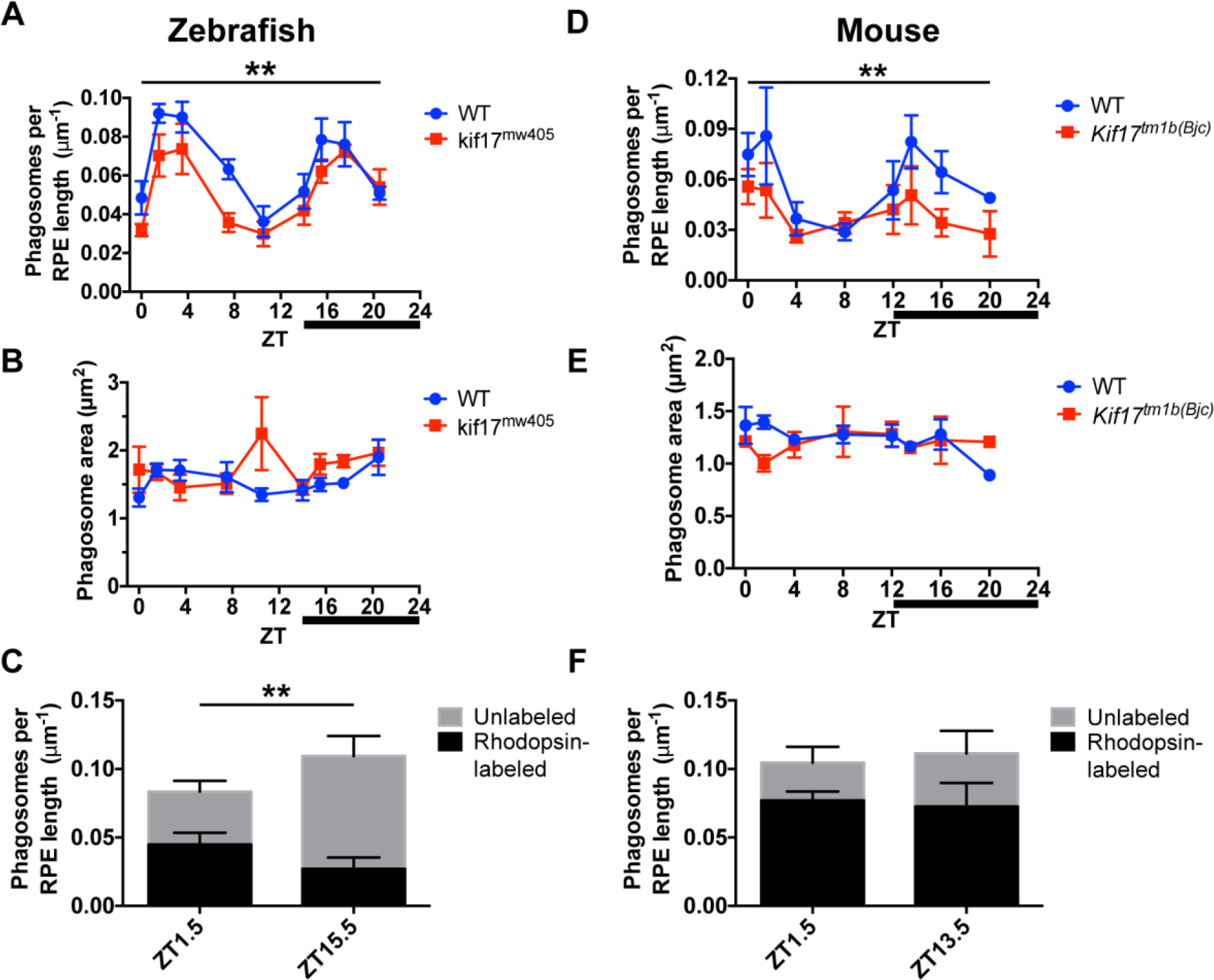
Loss of *kif17* diminishes disc shedding. **(A)** TEM analysis of phagosome number in 14 dpf wild-type (n=5 larvae at each timepoint, 1400 ± 97 μm of RPE measured at each timepoint) and *kif17*^*mw405*^ (n=5 larvae at each timepoint, 1469 ± 36 μm of RPE measured at each timepoint) zebrafish collected at nine discrete timepoints following light onset (ZT 0). Dark onset begins at ZT 14 (indicated by dark bar). Two-way ANOVA was performed to determine significance in both phagosome number rhythmicity throughout the timepoints (p<0.0001, ****) as well as between genotypes (p=0.0011, **). (B) TEM analysis of phagosome size in 14 dpf wild-type (n=5 larvae at each timepoint, 93 ± 14 phagosomes measured at each timepoint) and *kif17*^*mw405*^ (n=5 larvae at each timepoint, 79 ± 10 phagosomes measured at each timepoint) zebrafish. (C) Immunogold labeling of rhodopsin-containing phagosomes in 14 dpf wild-type retina at either the morning peak of disc shedding, ZT 1.5 (n=5 larvae, 1738 μm of RPE),or the evening peak, ZT 15.5 (n=5, 1341 μm of RPE). Two-way ANOVA was performed to determine significance in the interaction between the time of day and the concentration of rod or cone phagosomes (p=0.0082, **). There were no statistical differences in the total number of phagosomes between the morning or night peak. Bonferroni’s multiple comparisons post-hoc test was performed to determine significance in the increased of unlabeled phagosomes between morning and night (p=0.0158, *). (D) TEM analysis of phagosome number in 2-4 month wild-type (n=3 mice at each timepoint, 979 ± 42 μm of RPE measured at each timepoint) and *Kif1T^m1b(Bjc)^* (n=3 mice at each timepoint, 940 ± 45 μm of RPE measured at each timepoint) mice collected at eight discrete timepoints following light onset (ZT 0). Dark onset begins at ZT 12 (indicated by dark bar). Two-way ANOVA was performed to determine significance in both phagosome number rhythmicity throughout the timepoints (p=0.0300, *) as well as between genotypes (p=0.0096, **). (E) TEM analysis of phagosome size in 2-4 month wild-type (n=3 mice at each timepoint, 54 ± 7 phagosomes measured at each timepoint) and *Kif17^tm1b(Bc)^* (n=3 mice at each timepoint, 35 ± 4 phagosomes measured at each timepoint) mice. **(F)** Immunogold labeling of rhodopsin-containing phagosomes with B630 rhodopsin antibody in mouse retina at either the morning peak of disc shedding, ZT 1.5 (n=5 mice, 554 μm of RPE), or the evening peak, ZT 13.5 (n=5, 594 μm of RPE). Two-way ANOVA was performed to determine no significance differences in the interaction between the time of day and the concentration of rod or cone phagosomes (p=0.5700) or in the differences in phagosome number between the morning or night peak (p=0.8076), but there was a significant increase in the total number of rod phagosomes compared to cone phagosomes (p=0.0074, **).

To more carefully analyze the differences in disc shedding patterns between wild-type and *kif17*^*mw405*^ larvae as well as between the morning and evening peaks, we performed spline interpolation, a method of constructing a piecewise polynomial curve to smoothly fit the disc shedding data across all timepoints, for both the wild-type and *kif17*^*mw405*^ phagosome number data **(Figure S9A)**. For wild-type larvae, the two peaks in the number of phagosomes occurred 2.5 hours after light onset and subsequently 2 hours after dark onset **(Table S1)**. In comparison, *kif17*^*mw405*^ mutants had estimated peaks at 2.5 hours after light onset, and 3.2 hours after dark onset, suggesting that, in addition to the decrease in the total number of phagosomes shed associated with loss of *kif17*, the kinetics of disc shedding at night might also be altered. Additionally, the kinetics of phagosome digestion could be roughly estimated by the decreasing slope in the number of phagosomes observed following either the morning or night peak. However, this is under the assumption that there are no new phagosomes being shed during this period. Following the morning peak of disc shedding, we estimated similar slopes of 8x10^-3^ and 10X10^-3^ phagosomes/μm of RPE per hour in wild-type and *kif17*^*mw405*^ larvae respectively. Again, after the evening peak we estimated similar slopes of 9x10^-3^ and 8x10^-3^ phagosomes/μm of RPE per hour for wild-type and *kif17*^*mw405*^ larvae respectively. Taken together, these data suggest that the halflife of a phagosome of approximately 3-4 hours is unaffected by the loss of *kif17*.

In addition to the deficiencies in disc shedding caused by loss of *kif17* in zebrafish, we observed a similar diminution in total number of phagosomes in *Kif17* deficient mice **(Figure 5D)** without a significant impact on phagosome size **(Figure 5E)**. In contrast to the report that rat disc shedding occurs in only a single peak following light onset [21], we observed two peaks of disc shedding in mice, more comparable to our findings in zebrafish: one shortly after light onset and another shortly after dark onset. We further performed spline interpolation on both the wild-type and *Kif17*^*tm1b(Bjc)*^ phagosome number data **(Figure S9B)** and estimated that the morning peak in phagosome number occurred 1.4 hours after light onset in wild-type compared to 0.7 hours in *Kif17*^*tm1b(Bjc)*^ mice, while the evening peak was estimated to occur at about 1.7 hours after dark onset for both genotypes **(Table S2)**. Following the morning peak of disc shedding, phagosome number decreased at similar rates of 9x10^-3^ and 11x10^-3^ phagosomes/μm of RPE per hour in wild-type mice compared *Kif17*^*tm1b(Bjc)*^ mice. After the evening peak, the number of phagosomes decreased at similar rates of 6x10^-3^ and 7x10^-3^ phagosomes/μm of RPE per hour in wild-type and *Kif17*^*tm1b(Bjc)*^ mice respectively. Under all conditions, the half-life of a phagosome in mice is approximately 1-2 hours. As in zebrafish, this analysis suggests little if any alteration in the kinetics of phagosome accumulation or digestion from loss of *Kif17*. In both mice and zebrafish, loss of *Kif17* is associated with a decrease in the total number of phagosomes, supporting a role for KIF17 in promoting disc shedding in the mature photoreceptor.

It has been suggested in an analysis using goldfish [22] that morning and evening peaks reflect disc shedding by rods and cones respectively, although the two types of phagosomes cannot be readily distinguished after phagocytosis. To further define these two peaks of disc shedding in zebrafish, we performed the immunogold rhodopsin labeling of phagosomes in wild-type larvae at both the morning (ZT 1.5) and evening (ZT 15.5) peaks. While there was no change in the total number of phagosomes at either of these peaks, there was an increase in the number of unlabeled phagosomes at the evening peak, suggesting an increase in cone disc shedding **(Figure 5C)**. However, at least 25% of the phagosomes present at the night peak were still rod phagosomes. Additionally, we performed immunogold labeling with the cone-specific transducin-α antibody, and again found that while there was no change in the total number of phagosomes at either the morning or evening peak, the evening peak contains a higher ratio of cone to rod phagosomes than the morning peak **(Figure S8C)**. However, cone phagosomes still composed about 40% of the phagosomes in the morning peak, while unlabeled phagosomes compose about 40% of the evening peak. Overall, there are two peaks of phagosomes in zebrafish: one shortly after light onset with a slightly higher proportion of rod phagosomes and another peak occurring shortly after dark onset with a slightly higher proportion of cone phagosomes. These zebrafish data ultimately do not support the absolute distribution of cone shedding at night and rod shedding during the day as is commonly reported in the literature [22], but rather only a small bias of cone shedding at night and rod shedding during day. Additionally, we performed immunogold rhodopsin labeling of phagosomes in mice with B630, a mouse monoclonal anti-rhodopsin, at both the morning (ZT 1.5) and evening (ZT 13.5) peaks. At both peaks, the majority of phagosomes contained rhodopsin, although there was a small proportion of unlabeled phagosomes presumably from cones **(Figure 5F)**. Of note, there was no significant increase in the number of these unlabeled phagosomes at the night-time peak.

### Constitutively active CaMKII promotes disc shedding in a Kif17 dependent manner

While we have shown that amino acid changes at the conserved zebrafish S815 phosphorylation site can control KIF17 localization and promote photoreceptor disc shedding, we sought to associate this regulation more directly to CaMKII, which phosphorylates this conserved serine of KIF17 in neurons [11] and is expressed in cones [23]. We injected tCaMKII-GFP, a constitutively active form of CaMKII [24], under control of the TaCP cone promoter and investigated disc shedding at 7dpf as compared to control larvae injected with soluble GFP under control of the same cone promoter. We found that wild-type larvae injected with tCaMKII-GFP had a two-fold increase in phagosome number as compared to soluble GFP controls. However, this increase in disc shedding did not occur when tCaMKII-GFP was injected in *kit17*^*mw405*^ larvae **(Figure 6A)**. There was no change in phagosome size associated with any of the injected larvae **(Figure 6B)**. Ultimately, these data show that the effect of tCaMKII depends on the presence of endogenous Kif17 and implicates CaMKII mediated phosphorylation of Kif17 in the regulation of disc shedding.

**Figure 6.**
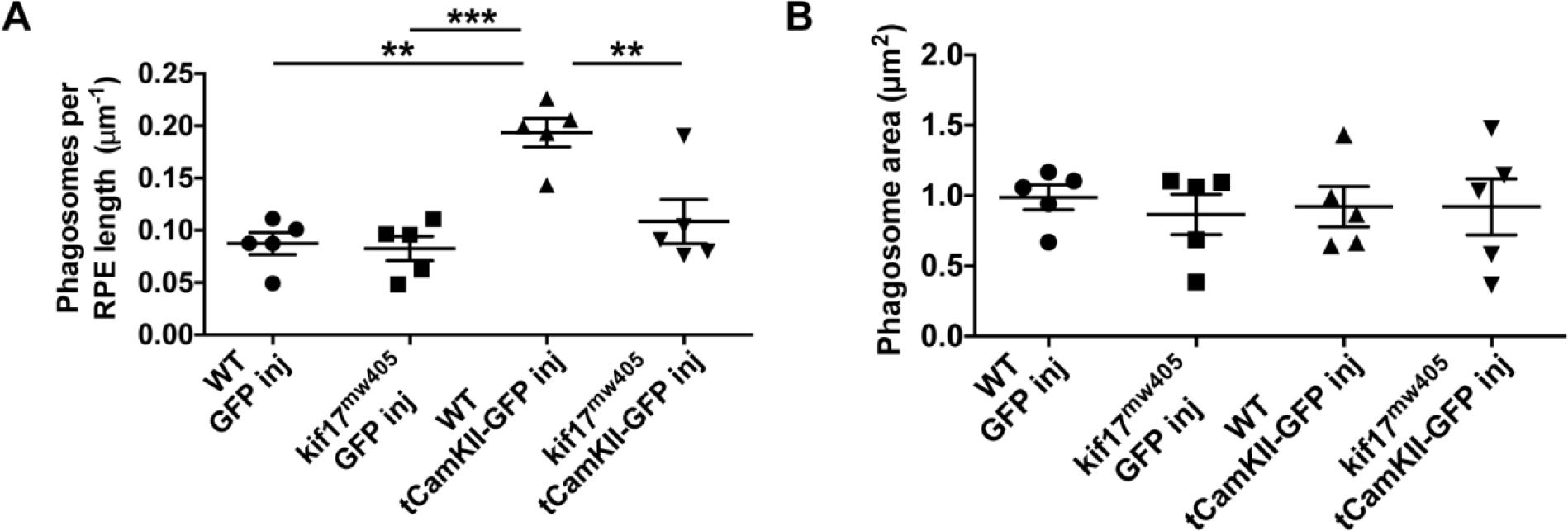
CaMKII promotes *kif17-dependent* disc shedding. **(A)** TEM phagosome number analysis of 7 dpf GFP-positive wild-type and *kif17^mw40S^* larvae injected with either control TaCP:GFP (n=5 injected larvae, 843 μm of RPE for WT; n=5, 795 μm of RPE for *kif17*^*mw405*^) or the constitutively active CaMKII construct TaCP:tCaMKII-GFP (n=5, 1011 μm of RPE for WT; n=5, 1322 μm of RPE for *kif17^mw405^)*. **(B)** Quantification of the size of phagosomes for wild-type and *kif17*^*mw405*^ larvae injected with either TaCP:GFP (n=5 injected larvae, 74 phagosomes for WT; n=5, 70 phagosomes for *kif17*^*mw405*^) or TaCP:tCaMKII-GFP (n=5, 193 phagosomes for WT; n=5, 141 phagosomes for *kif17^mw405^)*.

## Discussion

### Phosphorylation promotes ciliary localization of KIF17

In this work, we show that phospho-mimetic KIF17 has a strongly enhanced ciliary localization in four different mammalian cell lines and in zebrafish photoreceptors. We propose two distinct models for the regulation of ciliary localization of KIF17 via phosphorylation **(Figure 7)**. In the first model, phosphorylation of KIF17 via CaMKII could promote the ciliary entry of KIF17. However, because both wild-type and phospho-deficient KIF17 have a basal level of ciliary localization, ciliary entry of non-phosphorylated KIF17 does not appear to be entirely restricted. In a second model, phosphorylation could promote accumulation within the OS, potentially at the distal tip as has been shown previously [6, 7, 15-17], through an interaction with a plus-end microtubule associated protein. KIF17 has a consensus sequence for EB1-binding and microtubule plus-end accumulation [25] and has been shown to interact with EB1 in epithelial cells for regulating microtubule dynamics at the plus-ends of cytoplasmic microtubules [26]. EB1 also localizes to the axonemal tip of flagella in *Chlamydomonas reinhardtii* [27]. Thus, enhanced affinity for EB1 (or another related microtubule associated protein) via phosphorylation of KIF17 could lead to the increased ciliary localization. De-phosphorylation could subsequently release KIF17 from this interaction, leading to removal from the cilia via either active transport or diffusion. In support of this model, truncated KIF17 lacking the motor domain is still able to accumulate at the ciliary tip [7, 15, 17], implying the need for some additional microtubule-associated protein for KIF17 ciliary localization.

**Figure 7.**
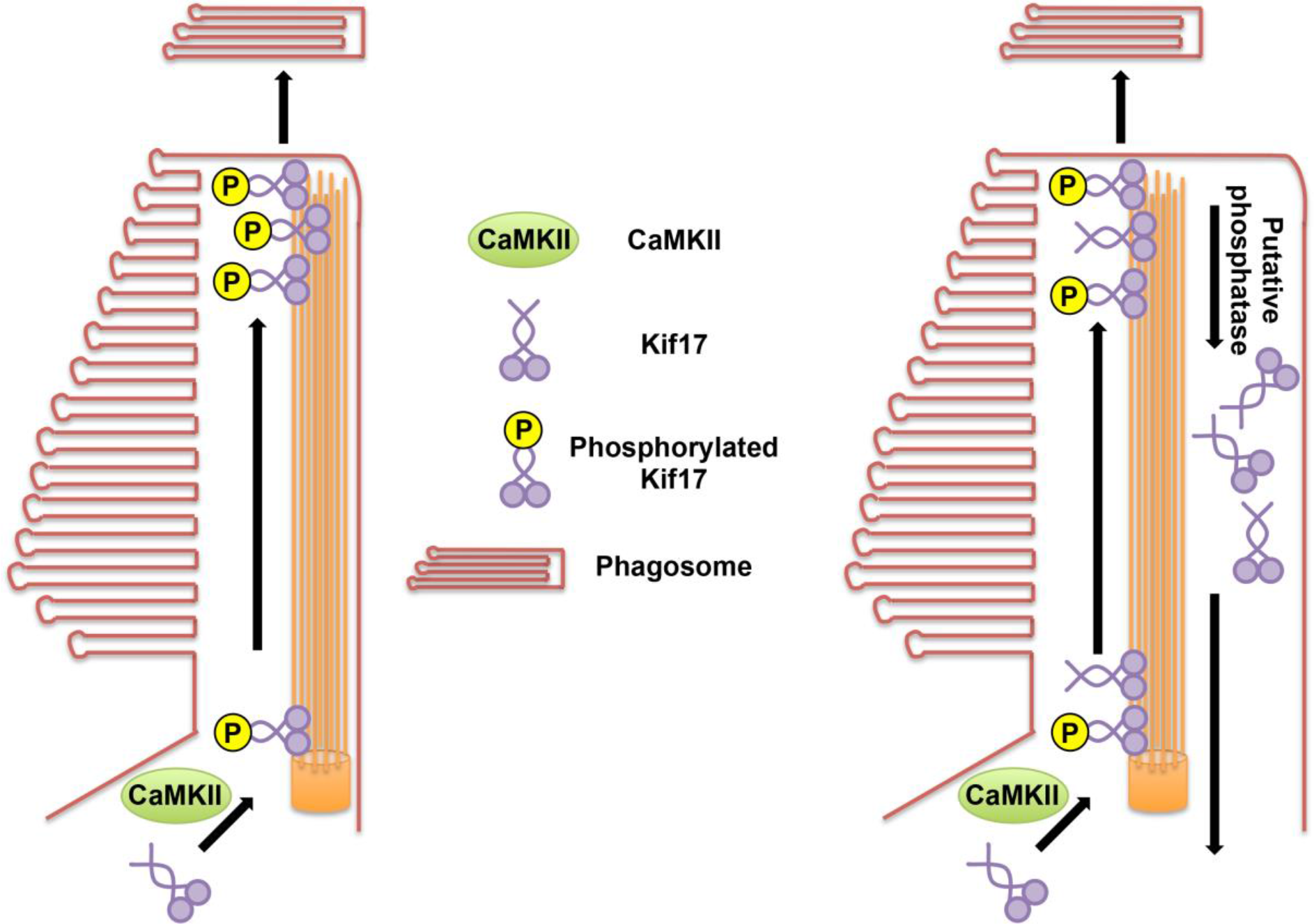
Models of regulation of Kif17 localization through phosphorylation. (Left) In the first model, ciliary localization of Kif17 could be controlled specifically at the transition zone, where only phosphorylated Kif17 (presumably mediated through CaMKII) would enter the cilium. In a photoreceptor, KIF17 in the OS promotes cell-autonomous disc shedding through some putative photoreceptor-centric signaling process. (Right) In a second model, ciliary localization of Kif17 could be controlled not at the transition zone, but rather specifically at the ciliary tip. Both phosphorylated and non-phosphorylated Kif17 could enter the cilium, but non-phosphorylated Kif17 only transiently associates with the axonemal tip. Phosphorylated Kif17 has an enhanced interaction with some putative microtubule associated protein at the axonemal tip, leading to an enhanced ciliary accumulation. A putative phosphatase could regulate Kif17 localization at the ciliary tip, and non-phosphorylated Kif17 could subsequently exit the cilium through either active or passive transport. Similar to the other model, accumulation of Kif17 at the distal tip of the photoreceptor OS promotes cell-autonomous disc shedding.

In contrast to the strong promotion of ciliary localization, there was only a mild, cell line-dependent restriction of nuclear localization associated with phospho-mimetic KIF17. There are two explanations for these data: that KIF17 phosphorylation can directly inhibit nuclear entry only in certain cellular contexts or that the mild effects on nuclear entry are secondary effects of the strong promotion of ciliary localization. However, because all four cell lines depict a strong promotion of ciliary localization of phospho-mimetic KIF17, it is unlikely that the effects on nuclear localization seen in only two of the cell lines are secondary effects. Ultimately, this evidence suggests that KIF17 phosphorylation is a robust regulator of ciliary localization in a variety of cell types, including photoreceptors, and in a variety of species, with much milder effects on nuclear localization.

### KIF17 positively regulates disc shedding

The function of Kif17 in zebrafish photoreceptors has been ambiguous. While early morpholino knockdown experiments suggested that Kif17 was important for OS development [12], two genetic mutants [28, 29] were shown to have morphologically normal OS. Similarly, mice were shown to have morphologically normal photoreceptors [15]. We recently resolved these apparent contradictions by showing that loss of Kif17 delays the onset of OS morphogenesis in both zebrafish and mice [14]. Nonetheless, OS in both species attain normal morphology and dimensions [14, 15], raising the question of whether Kif17 functions in a mature photoreceptor.

In this work, we find significant decreases in disc shedding in both *kif17*^*mw405*^ zebrafish and *Kif17*^*tm1b(Bjc)*^ mice. While disc shedding is not completely ablated with loss of *Kif17*, we have previously proposed that additional factors, such as kinesin-II or diffusion, can compensate for defects in trafficking in *Kif17* mutant photoreceptors [14]. Whether other factors compensate for loss of KIF17’s function in promoting disc shedding and whether attenuating such factors can lead to compound effects in *Kif17* mutants on disc shedding remains to be elucidated. Additionally, cone expression of transgenic phospho-mimetic Kif17 results in a cell-autonomous enhancement of disc shedding. CaMKII appears to be one of the kinases responsible for phosphorylation of Kif17 in photoreceptors *in vivo*, as cone expression of constitutively active tCaMKII promotes disc shedding in a *kif17*-dependent manner. However, additional investigation is required to identify other potential kinases or the phosphatase that may regulate removal of Kif17 from the OS. Ultimately, the phospho-mimetic Kif17 “gain of function” effects in promoting disc shedding combined with decreased disc shedding phenotype associated with genetic loss of Kif17 strongly implicate Kif17 in an important role regulating disc shedding at the level of the photoreceptor. Major advances have been made regarding the regulation of disc shedding, but most of that work has been at the level of phagocytosis by adjacent RPE cells [30-32]. However, PS exposure at the distal OS precedes shedding events [19], and these data on Kif17 further imply an active process in the ciliary OS tip that promotes detachment of the distal domain and subsequent phagocytosis.

Another significant conclusion from this work is that photoreceptors can exhibit considerable plasticity in the kinetics of ciliary OS turnover while still maintaining normal dimensions. It is well known that mouse rods shed approximately 10% of their length daily and maintain a steady-state length through compensating OS renewal [33]. In the *Kif17* mutant mice and zebrafish, diminished disc shedding is not accompanied by reduced size of the OS [14], suggesting an overall slower rate of renewal at the OS base. Likewise, the dimensions of the OS are unchanged in the “gain of function” experiments involving phospho-mimetic transgene expression, implying that enhanced disc shedding is compensated for by enhanced OS formation. While we propose that Kif17 is implicated in regulating disc shedding directly given this work, it is possible that Kif17 is involved in promoting new morphogenesis at the OS base and the increased disc shedding is a secondary effect to maintain a steady-state OS. Although the open disc conformation of cone photoreceptors complicates experiments, techniques involving autoradiography [34] and fluorescent [35] pulse-labels to study the dynamics of rod OS morphogenesis can be applied to investigate the changes in rates of new cone OS morphogenesis associated with either gain or loss of function of Kif17.

As little is known about the photoreceptor-derived signaling that regulates disc shedding, the specific mechanism by which Kif17 promotes disc shedding is unclear. However, as PS exposure at the distal OS precedes disc shedding [19], it is possible that Kif17 is involved in regulating the localization of a putative scramblase [36] or floppase [37] involved in mediating PS transport from the cytosolic to extracellular leaflet of the OS membrane. Alternatively, Kif17 accumulated at the distal OS could interact with various ligands expressed in the photoreceptor, such as SEMA4D, that are involved in regulating disc shedding via receptors in the RPE [38]. Future work will address potential interacting partners of Kif17 to reveal how accumulation at the distal OS promotes disc shedding.

### Temporal regulation of cone and rod disc shedding

Our investigation of Kif17 loss of function required analysis of disc shedding at multiple timepoints because photoreceptor turnover is well-known to occur on a diurnal schedule [21, 22, 39], which led to unexpected surprises in our analysis in both zebrafish and mice. First, it is generally thought that rods shed in the morning after light onset, while cones do so in the evening after dark onset [22, 40, 41]. However, this has been generally determined by counting phagosomes in the RPE where the differences between the two cell types are not apparent. Our new data using immunogold labeling specific for either rods (using the rhodopsin antibody) or cones (using the GNAT2 antibody) shows that both rod and cone phagosomes are present at both day-time and night-time peaks in zebrafish. Our interpretation of these data is that both rods and cones shed discs at either time of day, although there is a slight preponderance of rod phagosomes in the morning and cones at night. At the very least, the absolute idea of “rods shed during the day and cones at night” [22] must be reconsidered. Secondly, we observed equal peaks of disc shedding in mice both after light onset and offset, which contrasts with the long-held view that disc shedding in rod-dominated rodents occurs principally after light onset [21]. This can hardly be explained as a dichotomy of rod versus cone shedding as the mouse retina consists of ~97% rods and ~3% cones [42]. A key difference in our analysis in mice is that we sampled at symmetric time points, including both 1.5 hours after both light onset and dark onset, whereas other studies have used wider sampling intervals such as 4 hour intervals, possibly missing important events occurring after dark onset [21]. This surprising finding is of potential broad significance in understanding disc shedding and OS turnover and is under current investigation.

## Conclusion

Overall, we show that phosphorylation of KIF17 promotes its ciliary localization. In cone photoreceptor outer segments, phosphorylated KIF17 promotes cell-autonomous disc shedding. As disc shedding has been predominantly studied within the retinal pigment epithelium, this work implicates photoreceptor-derived signaling in the underlying mechanisms of disc shedding.

## Methods

### Mouse and zebrafish husbandry

C57BL/6 mice were maintained under 12 hour:12 hour light:dark cycle. We generated the *Kif17*^*tm1b(Bjc)*^ mouse line as described previously [14]. Adult ZDR zebrafish were housed at 28.5°C on a 14 hour:10 hour light:dark cycle. Embryos were raised at 28.5°C on a 14 hour:10 hour light:dark cycle. We generated the *kif17*^*mw405*^ zebrafish line as described previously [14]. All experiments were approved and conducted in accordance with the Institutional Animal Care and Use Committee of the Medical College of Wisconsin. For all experiments involving animals, a biological replicate is defined as a single mouse or single zebrafish. Male and female animals were randomly assigned to each experimental group. For both zebrafish and mice, littermate animals were used.

### Cloning of constructs

A plasmid encoding CMV:KIF17-mCherry was generated as previously reported [16]. Overlap extension PCR was performed to create the two mouse phospho-mutants: CMV:KIF17(S1029D)-mCherry and CMV:KIF17(S1029A)-mCherry. The wild-type codon sequence for S1029, AGC, was mutated to GAC for S1029D and GCC for S1029A. A plasmid encoding TaCP:Kif17-GFP was generated as previously reported [16]. Overlap extension PCR was performed to create the two zebrafish phospho-mutants: TaCP:Kif17(S815D)-GFP and TaCP:Kif17(S815A)-GFP. The wild-type codon sequence for S815, AGC, was mutated to GAC for S815D and GCC for S815A. The constitutively active CaMKII (tCaMKII-GFP) was generated by Yasunori Hayashi [24]. tCaMKII-GFP was cloned downstream of the TaCP promoter using the Gateway Cloning system (Invitrogen). The control TaCP:GFP plasmid was generated by Breandán Kennedy [43].

### Zebrafish transgenesis

For various experiments involving injected larvae, 9.2nL of a working solution containing 25ng/μL of a plasmid (encoding either TaCP:Kif17-GFP, TaCP:Kif17(S815D)-GFP, TaCP:Kif17(S815A)-GFP, TaCP:tCaMKII-GFP, or TaCP:GFP), as well as 0.05% phenol red, were injected into ZDR embryos at the 1-cell stage. For stable line generation of TaCP:Kif17-GFP, TaCP:Kif17(S815D)-GFP, and TaCP:Kif17(S815A)-GFP, 10ng/μL transposase RNA was included in the injection solution to facilitate genome integration [44]. Injected embryos were subsequently raised and outcrossed to screen for germ-line transmission. GFP-positive embryos were raised and outcrossed for several generations to establish a stable line of low-expressing, single insert transgenic fish to minimize non-specific effects due to high levels of over-expression **(Figure S5A,B)**. For experiments, embryos were imaged with epifluorescence at 4dpf and fluorescence intensity of eyes was quantified to ensure an identical expression level between the three transgenes **(Figure S5A,C)**.

### Antibodies

Commercial primary antibodies included: 6-11B-1, or mouse monoclonal anti-acetylated α-tubulin (Sigma), rabbit polyclonal anti-GNAT2 (MBL International). The rabbit polyclonal anti-green and blue cone opsin antibodies were gifts from Dr. Thomas Vihtelic (University of Notre Dame) [45]. The mouse monoclonal anti-rhodopsin antibodies K62-171c and B630 were gifts from Dr. Paul Hargrave (University of Florida) [20]. Nuclei were stained with Hoechst (Thermo Scientific). Fluorescent secondary antibodies included goat anti-mouse IgG Alexa Fluor 488 and goat antirabbit IgG Alexa Fluor 594. Immunogold labeling secondary was either goat anti-mouse IgG 10nm colloidal gold (Electron Microscopy Sciences) or goat anti-rabbit IgG 10nm colloidal gold (Electron Microscopy Sciences).

### Mammalian cell culture analysis

LLC-PK1 (ATCC CL-101), HEK-293 (ATCC CRL-1573), hTERT-RPE1 (ATCC CRL-4000), and IMCD3 (ATCC CRL-2123) were obtained from ATCC. Cell line authentication and testing of mycoplasma-negative were performed by ATCC. Additional testing of mycoplasma infection is performed biannually with MycoAlert (Lonza). LLC-PK1 cells were grown in Medium 199 (Gibco), HEK-293 cells were grown in DMEM (Gibco), and hTERT-RPE1 and IMCD3 cells were grown in DMEM/F12 (1:1) (Gibco). All media were supplemented with 10% FBS and 1% penicillin and streptomycin and cells were grown at 37°C in 5% CO_2_. For ciliogenesis and transfection, cells were transferred to Opti-MEM (Gibco) at 60-80% confluency and transiently transfected with Lipofectamine 3000 (Invitrogen). Serum-starvation in Opti-MEM induced ciliogenesis and after 24 hours, cells were fixed in 4% PFA for 20 minutes and subsequently immunostained with acetylated a-tubulin to indicate cilia and Hoechst to label nuclei. Because of the nature of transient transfection, there were a variety of transgenic expression levels. Approximately 20% of cells with a high level of expression in which transgene appeared to be highly vesicular or stuck in Golgi were excluded from analysis. For analysis, five separate transfections of each transgene in each cell line were performed. For ciliary localization in each cell line, the number of cells with mCherry labeling of the cilia was counted and divided by the total number of transfected cells. For nuclear localization, ImageJ was used to quantify the average intensity of both cytoplasmic and nuclear mCherry signal. Following subtraction of the average intensity of the background of each image (quantified from an area where there was no cellular fluorescence), the ratio of the cytoplasmic to nuclear average intensity was calculated.

### Photoreceptor localization analysis

Embryos were injected with constructs as described above for mosaic expression and raised in phenylthiourea (PTU) to delay melanin pigment formation. At 4 dpf, larvae were anesthetized with tricaine and screened for GFP fluorescence. Immunofluorescence staining was performed on cryosections of 5 dpf GFP-positive larvae as previously described [12] with antibodies to green or blue opsin to label cone OS and Hoechst to label nuclei. For quantification, only injected larvae with positive transgenic expression were analyzed. For OS localization, GFP labeling in the opsin-labeled OS was analyzed compared to the total number of transgene-expressing cones. For line intensity analysis, a line along the length of the OS (as determined by opsin-labeling) was used to calculate GFP signal intensity in ImageJ. For standardization of levels of transgene expression, all intensity values along a single line were standardized to the maximum value along that line. Additionally, to standardize for variations in lengths of the OS, the length of an OS was split into ten equal groups based on position from base to distal OS and intensity values were averaged for each of these ten groups to generate data for a single technical replicate. In each biological replicate, several technical replicates were averaged, as described in captions.

### Disc shedding analysis

For transgenic or injected larva disc shedding analyses, zebrafish embryos were raised in phenylthiourea until 4 dpf when they could be screened for GFP fluorescence. GFP-positive embryos were then raised in regular fish water until either 5dpf **(Figure S4)** or 7dpf (rest of the experiments), when they were collected at 1.5 hours following light onset. For zebrafish diurnal disc shedding analysis, 14dpf wild-type and *kif17*^*mw405*^ larvae were collected at nine different timepoints following light onset: 0 (light onset), 1.5 hours, 3.5 hours, 7.5 hours, 10.5 hours, 14 hours (dark), 15.5 hours, 17.5 hours, 20.5 hours. Collected larvae were processed as previously described for TEM [14]. For mouse diurnal disc shedding analysis, 2- to 4-month old C57BL/6 and *Kif17*^*tm1b(Bjc)*^ litter-mate mice were collected at eight different timepoints following light onset: 0 (light onset), 1.5 hours, 4.0 hours, 8.0 hours, 12 hours (dark), 13.5 hours, 16 hours, and 20 hours. Mice were euthanized and whole eyes were processed as previously described for TEM [14]. For zebrafish analysis, five larvae were analyzed for each genotype and timepoint for a total of 45 larvae per genotype. For mouse analysis, three mice were analyzed for each genotype and timepoint for a total of 24 mice per genotype. For phagosome number analysis, the number of phagosomes was counted and divided by the total length of RPE measured. For phagosome size analysis, phagosome area was measured and averaged for each sample. For OS size analysis, OS area was measured and averaged for each sample.

### Spline interpolation analysis

For zebrafish, cubic spline interpolation was performed in Prism 6 (GraphPad) with 40 segments over the 24 hour period (including the ZT 0 timepoint repeated at ZT 24) for increments of 0.615385 hours. For mice, cubic spline interpolation was performed with 36 segments for increments of 0. 6857143 hours. For analysis, the maximum disc shedding rate during both the morning and evening was determined as the peak of disc shedding. To calculate the digestion rates of phagosomes, the maximum and minimum points of disc shedding for both the morning and evening shedding events was determined. Rates of phagosome digestion were subsequently calculated as the decreasing slope of the number of phagosomes over the period of between 25% and 75% of the total time between maximum and minimum. Phagosome half-life was calculated for both morning and evening as the time after the maximum peak until the phagosome number reached half-way between the maximum and minimum points.

### Immunogold labeling

For immunogold analysis, 7dpf transgenic larvae were collected at 1.5 hours after light onset or 14dpf wild-type larvae were collected at both 1.5 hours after light onset and 1.5 hours after dark onset. Larvae were fixed overnight in 0.1% glutaraldehyde and 2% paraformaldehyde in 0.1M phosphate buffer (pH 7.4). Larvae were washed in phosphate buffer, dehydrated with two 30 minute washes of 70% ethanol, and incubated with a 2:1 mixture of LR white:70% ethanol for 1 hour. Larvae were then incubated overnight in LR white at room temperature. In the morning, following two 30 minute changes of LR white, larvae were embedded and polymerized at 55°C for 48 hours. Ultra-thin sections were mounted on 200 mesh formvar/carbon coated copper grids. Grids were incubated on drops of 0.15 mM glycine for 10 minutes, then on 5% BSA in 0.1M phosphate buffer for 15 minutes. For labeling, grids were incubated on drops of antibody for 90 minutes. Grids were washed in 5% BSA in 0.1M phosphate buffer and incubated with colloidal gold secondary (Electron Microscopy Sciences) for 1 hour. Grids were washed in double-distilled H2O and stained with 1% aqueous uranyl acetate for 2 minutes. For each analysis, a total of five larvae were analyzed for each condition. For analysis, the number of both gold-labeled and unlabeled phagosomes was counted and divided by the total length of RPE measured.

### qPCR analysis

For mouse circadian *Kif17* expression analysis, adult C57BL/6 mouse retinas were collected every four hours beginning at light onset over a 24 hour period and placed RNAlater (Ambion) overnight at 4°C. For zebrafish circadian *kif17* expression analysis, 14dpf ZDR zebrafish eyes were collected at nine discrete timepoints following light onset: 0 (light onset), 1.5 hours, 3.5 hours, 7.5 hours, 10.5 hours, 14 hours (dark), 15.5 hours, 17.5 hours, 20.5 hours. For each biological replicate, three zebrafish eyes were pooled. For zebrafish opsin expression analysis, five whole 7dpf zebrafish larvae were pooled. Following sample collection, RNA was extracted, cDNA synthesized, and qPCR run as previously described [14]. For each data-point, three biological replicates were run in triplicate for both mouse and zebrafish samples. For mouse, published *Kif17* primers were used [46] and published RNA polymerase II primers were used as a reference gene [47]. For zebrafish, published ef1*α* primers were used as a reference gene [14], published opsin primers were used [14], and *kif17* primers are depicted below.

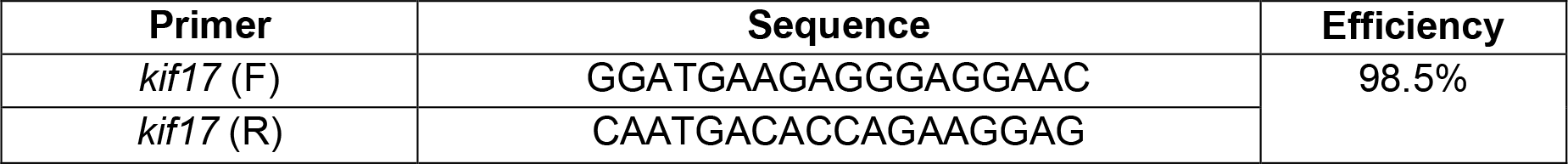

All expression values were standardized to 1 for the light onset timepoint (ZT 0).

### Imaging

Immunofluorescent images were acquired with a 100x, 1.30 NA oil-immersion lens (Nikon) on either an Eclipse TE300 (Nikon) microscope operating NIS-Elements (Nikon) and a Zyla sCMOS camera (Andor) or a C1 Plus-EX3 AOM Confocal System (Nikon) operating EZ-C1 (Nikon). TEM imaging was performed on a Hitachi H-600 with an ORCA-100 digital camera (Hamamatsu). Blinded image analysis was performed on de-identified, randomized images with ImageJ.

### Statistical analysis

For ciliary and nuclear localization of KIF17 in mammalian cell culture experiments, a one-way ANOVA was performed. Following significance, a Bonferroni multiple-comparisons post-hoc test was performed to determine significance between groups. For OS localization frequency and nuclear localization in zebrafish photoreceptors, a one-way ANOVA was performed. For line intensity analysis, a two-way ANOVA was performed. For single-timepoint disc shedding analysis in transgenic or injected larvae, a one-way ANOVA was performed. For immunogold phagosomes analysis in either transgenic or wild-type larvae, a two-way ANOVA was performed. For circadian disc shedding analysis in either zebrafish or mice, a two-way ANOVA was performed. For circadian *Kif17* mRNA expression in either mouse or zebrafish retinas, a one-way ANOVA was performed. For qPCR analysis of opsin expression, a one-way ANOVA was performed for each opsin gene. For all experiments, following significance with either one-way or two-way ANOVA, a Bonferroni multiple-comparisons post-hoc test was performed to determine significance between groups. For all experiments, values are expressed as mean ± SEM and significance is labeled as * : p ≤ .05, **: p ≤ .01, ***: p ≤ .001, ****: p ≤ .0001. Non-significant values are unlabeled.

## Abbreviations

RPE: retinal pigment epithelium
NLS: nuclear localization signal
CLS: ciliary localization signal
OS: outer segment
PS: phosphatidylserine
ZT: Zeitgeber Time

## Declarations

### Ethics approval and consent to participate

All experiments were approved and conducted in accordance with the Institutional Animal Care and Use Committee of the Medical College of Wisconsin.

### Consent for publication

Not applicable

### Availability of data and materials

All data generated or analysed during this study are included in this published article and its supplementary information files.

### Authors’ contributions

TRL, CI, and JCB designed experiments. TRL, CI, and SRK, performed experiments. TRL, BAL, and JCB analyzed data. TRL and JCB wrote the initial manuscript. All authors read and approved the final manuscript.

## Acknowledgements

Not applicable

